# Evaluation of the effects of nicotine on the friction of the articular cartilage and the reversion of the estrogen in vitro

**DOI:** 10.1101/2021.12.17.473251

**Authors:** Huiqin Li, Jiaxin Tang, Ying Zhang, Yao Han

**Affiliations:** Instrumental Analysis Center of Shanghai Jiao Tong University, Shanghai; Department of Orthopedics, Shanghai Fourth People’s Hospital Affiliated to Tongji University School of Medicine

## Abstract

**Background:** Osteoarthritis is a severe disease for menopausal women, especially for those who exposed in the smoking and second hand-smoking. This study investigated the effect of the nicotine and estrogen on the articular cartilage.

**Methods:** The articular cartilages were treated by nicotine and estrogen in vitro. Then the frictional properties and morphology on the surface were investigated using atomic force microscope. Proteoglycan 4(PRG4), as the key boundary lubricant of articular cartilage was characterized.

**Results:** Nicotine down-regulates the friction coefficient and secretion of PRG4 significantly and then the estrogen increase them again. The adhesion forces also showed the same trend due to the content of anti-adhesive PRG4.

**Discussion:** This study demonstrated that the present concentration nicotine has a negative effect on the articular cartilage and the estrogen has a better protecting effect. This may provide a potential guide for OA prevention and treatment.

## 1. Introduction

Osteoarthritis (OA) is a chronic joint disease characterized by progressive and irreversible degeneration of articular cartilage. Epidemiological investigation of OA showed that it is more severe in menopausal women, which may attribute to decreased levels of estrogen (Calvo et al., 2007; Christgau et al., 2004; Hoegh-Andersen et al., 2004; Mouritzen et al., 2003).

Smoking and second-hand smoking are important environmental risk factors of OA. Among numerous chemicals present in cigarettes, nicotine is one of the most physiologically active molecule (Kawakita et al., 2008). Previous studies indicated that nicotine had adverse effect on the proliferation and chondrogenic differentiation of Human Wharton’s jelly stem cells (hWJ-MSCs) which have a high proliferation rate and capability of differentiating into osteoblasts (Ruiz et al., 2012; Yang et al., 2017). Nicotine can delay skeletal growth(Deng et al., 2013; Kawakita et al., 2008) and bone healing (Krannitz et al., 2009). Moreover, smoking delays chondrogenesis in a mouse model of fracture healing(El-Zawawy et al., 2006). The secretome analysis of human articular chondrocytes suggested a negative effect of nicotine on the joint (Lourido et al., 2016). Our previous study showed that the nicotine can induce inflammation on articular cartilage (Peng et al., 2019). On the other hand, the effects of nicotine on the joint and the mechanisms involved in cartilage degradation remains controversial(Turner et al., 1997).

OA onset is associated with a loss of small and large proteoglycans(Poole et al., 1996). Local changes in perlecan expression and content have also been reported during OA development(Tesche and Miosge, 2004). Similar changes during OA have also been described specifically in the superficial zone, it shows heterogeneous expression(Felka et al., 2016).

Articular interfaces exhibit remarkable tribological function with synovial fluid. Osteoarthritis leads to an alteration in the composition of the synovial fluid, which is associated with an increase in friction and the progressive and irreversible destruction of the articular cartilage(Forsey et al., 2006; Morgese et al., 2017). Lubricin plays a pivotal role in joint boundary mode lubrication and the prevention of osteoarthritis(Jay et al., 2010; Jay and Waller, 2014; Young et al., 2006).

The relations between nicotine and estrogen and surface friction are not well defined. To gain a better understanding of the roles of nicotine on the joint, we investigated the effect of nicotine on friction and PRG4 secreted by surface zone articular chondrocytes at cell-level in this article. We found that nicotine down-regulates the friction coefficient and secretion of PRG4. On the other side, considerable efforts have been made to understand a better protecting effect of estrogen on cartilage. We also evaluated the impact of the estrogen, which can be help to understand the effect of smoking on menopausal women.

## 2. Materials and methods

### 2.1 Materials

Nicotine, Estrogen (17β-estradiol), Tamoxifen and Collagenase II were purchased from Sigma (St Louis, MO, USA). Dulbecco’ s modified Eagle’ s medium (DMEM), fetal bovine serum (FBS) and penicillin and streptomycin (Pen-Strep) were purchased from Gibco (New York, NY, USA). Trypsin (AR1007) was acquired from BOSTER (Wuhan, HB, CHN). 25 cm2 Cell culture flasks and 24-well plates were purchased from Corning (New York, NY, USA). Lactate dehydrogenase (LDH) Cytotoxicity Assay kit and DAPI were purchased from Beyotime (Shanghai, CHN).

### 2.2 Cell culture

Under aseptic conditions, chondrocytes were isolated from the superficial cartilage (approximately 100-150 μm thickness) of knee joints of 8 weeks old male New Zealand white rabbits. The animals were acquired from Experimental Animal Center of Shanghai Jiaotong University and were sacrificed with the approval of the Bioethics Committee of Shanghai Jiao Tong University.

The cartilage was minced into 0.1-0.2 mm tissue blocks and digested with 0.1% trypsin in phosphate buffer solution (PBS) in 5% CO_2_ at 37 °C for 30 min, followed by 0.02% collagenase II in high-glucose DMEM supplemented with 10% FBS and 1% Pen-Strep at 37 °C and 5 % CO_2_ for 20 hours. The digested tissue was filtered through a sterile 100-μm nylon sieve to remove debris and undigested tissue and the filtrate centrifuged at 1200 rpm for 8 min. Extracted chondrocytes were then cultured in sterile 25 cm^2^ cell culture flasks in high-glucose DMEM with 10% PBS and 1% Pen-Strep in 5% CO_2_ at 37 °C. The medium was changed every 3 days. Confluent cells were detached with 0.25% trypsin in PBS and split 1:2. Passage 2 chondrocytes were used for the current study.

### 2.3 *In vitro* exposure to nicotine and estrogen

All medium was changed to serum-free medium 12 hours before treatment. Chondrocytes were exposed to 8 different treatment groups (12 wells per group): (1) medium only; (2) medium + nicotine (0.06 mM); (3) medium + estrogen (1 nM); (4) medium + estrogen (1 nM); (5) medium + Tamoxifen (1 μM); (6) medium + Tamoxifen (1 μM) + nicotine (0.06 mM); (7) medium + Tamoxifen (1 μM) + estrogen (1 nM); (8) medium + Tamoxifen (1 μM) + nicotine (0.06 mM) + estrogen (1 nM); The concentration of nicotine, estrogen and Tamoxifen were extrapolated by previous studies (Baxter et al., 2018; Liang et al., 2017; Liu et al., 2018; Wang et al., 2004). After cultured in 5% CO_2_ at 37 °C, cytotoxicity assay, immunofluorescence and atomic force microscopy (AFM) were performed on 3 well per group.

### 2.4 *In vitro* immunofluorescence evaluation

For Sox 9, Collagen type II, Aggrecan and Lubricin immunofluorescence evaluation, cells were fixed in 4% (v/v) paraformaldehyde solution for 15 min and 0.5% (v/v) Triton X-100 in PBS for 40 min. Cells were rinsed 3 times with PBS for 5 min, and then treated with a blocking solution (5% bovine serum albumin in PBS) for 1h. The cells were incubated with primary antibodies diluted in blocking solution overnight at 4 °C. After rinsing 3 times in PBS for 5 min, the specimens were incubated with second antibody at room temperature for 1 hour. Cell nuclei were visualized by DAPI and cytoskeletons were detected by actin using TRITC-conjugated Phalloidin. Fluorescent images were captured under a confocal microscope system (Leica, TCS SP8 STED 3X, Germany). Fluorescence intensity and area was calculated using FIJI/ImageJ.

### 2.5 *In vitro* friction, adhesion, and surface roughness measurements evaluation via AFM

Nanoscale surface roughness measurements were acquired with an AFM (Dimension Icon, Bruker) in liquid cell with triangular silicon tips (DNP-E, Bruker Instruments, Santa Barbara, CA) of 20 nm nominal radius and 0.1 N/m spring constant. Surface scanning of 30μm*30μm and 10μm *10μm areas imaged, operating in PeakForce Tapping mode.

Friction, adhesion was measured with colloid cantilevers. PS particles with a diameter of 5μm were glued to the end of the cantilever(MLCT, tipless cantilevers, Bruker Instruments, Santa Barbara, CA) with the help of two-component adhesive(Araldite, Switzerland). The exact values of the normal and torsional spring constants were obtained by the thermal noise method(Green et al., 2004; Sader et al., 1999). The spring constant calibrations were carried out at a temperature of about 23 °C before particle attachment. All surface force measurements were performed at a constant approach and retraction velocity of 400 nm/s. At this velocity, the effects of hydrodynamic forces can be considered negligible(Thormann et al., 2008; Wang et al., 2013).

### 2.6 Statistical analysis

Data were expressed as mean ± standard deviation (SD). The differences between the groups were assessed using one-way analysis of variance (ANOVA) followed by post hoc Dunn-Bonferroni test with IBM SPSS Statistics for Windows, version 23.0 (IBM Corporation, Armonk, NY). P values <0.05 were considered statistically significant.

### 2.7 Approval statement

All animal experiments were carried out in compliance with the Guide for the Care and Use of Laboratory Animals of the Ministry of Health of China and NIH and the Declaration of Helsinki. The studies have been approved by the Bioethics Committee of Shanghai Jiao Tong University, China

## 3. Results

### 3.1 Friction coefficient *in vitro*

For the chondrocytes in surface zone articular cartilage, both of the nicotine treatments showed a general trend of higher friction compared to blank(M1) and adding estrogen <u>receptor antagonist</u>(M2) (Fig. 1). Nicotine increased μ of both M1 and M2 samples most significantly (μ_C_=0.15±0.02 vs μ_N_ =1.2±0.02 for M1 (P< 0.001) and μ_C_=0.3±0.02 vs μ_N_=2.54± 0.02 for M2 (P< 0.001). Estrogen treatments after nicotine treatment reduced μ of both M1 and M2 samples most significantly again (μ_N_=1.2±0.02 vs μ_NE_=0.6±0.01 for M1 (P< 0.001) and μ_N_=2.54± 0.02 vs μ_C_=1.24±0.02 for M2 (P< 0.0001)). Estrogen treatments only did not show significant differences both for M1 and M2.

**Fig.1.**
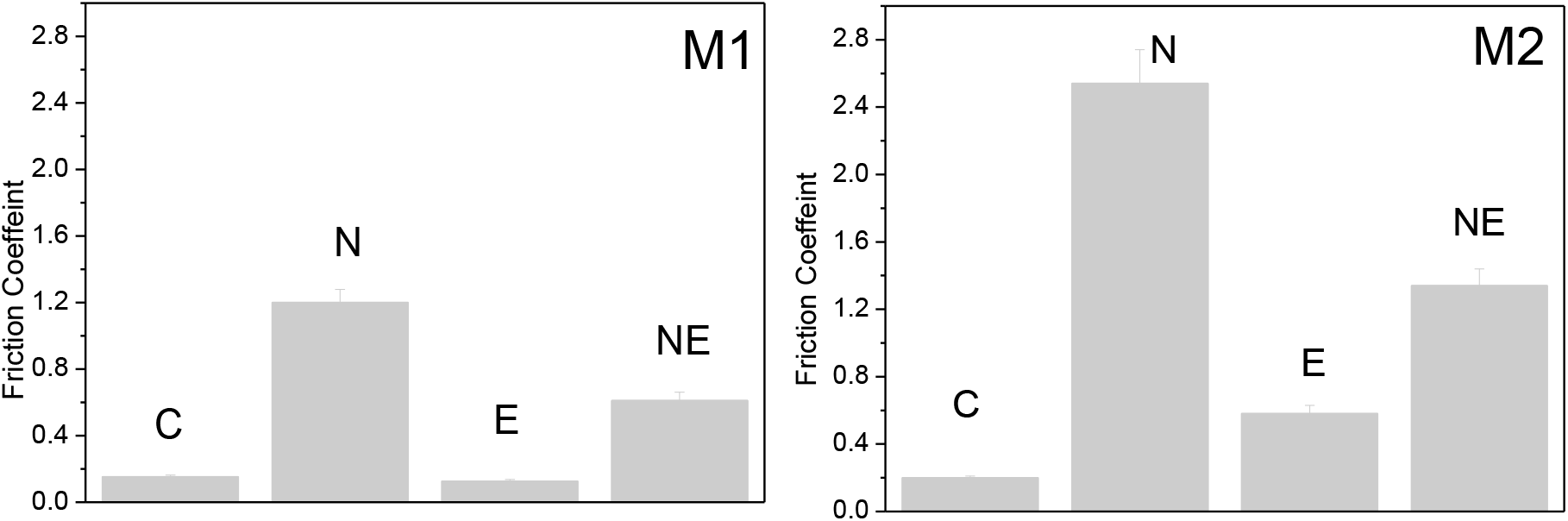
The friction coefficient of articular chondrocytes surfaces measured with AFM(M1 for normal chondrocytes, M2 for chondrocytes of adding estrogen receptor antagonist, C: control; N: nicotine treatment; E:only estrogen treatment; NE: estrogen treating after nicotine treatment) in vitro

### 3.2 Images and roughness *in vitro*

Surface roughness did not demonstrate a consistent trend or significant change for any nicotine and estrogen treatments. Topography images did not reveal any discernible differences between control and corresponding treated M1 and M2 sample (Fig.2). Fibrillated and amorphous surface structures were observed in the AFM images, in agreement with reports of previous studies.

**Fig.2.**
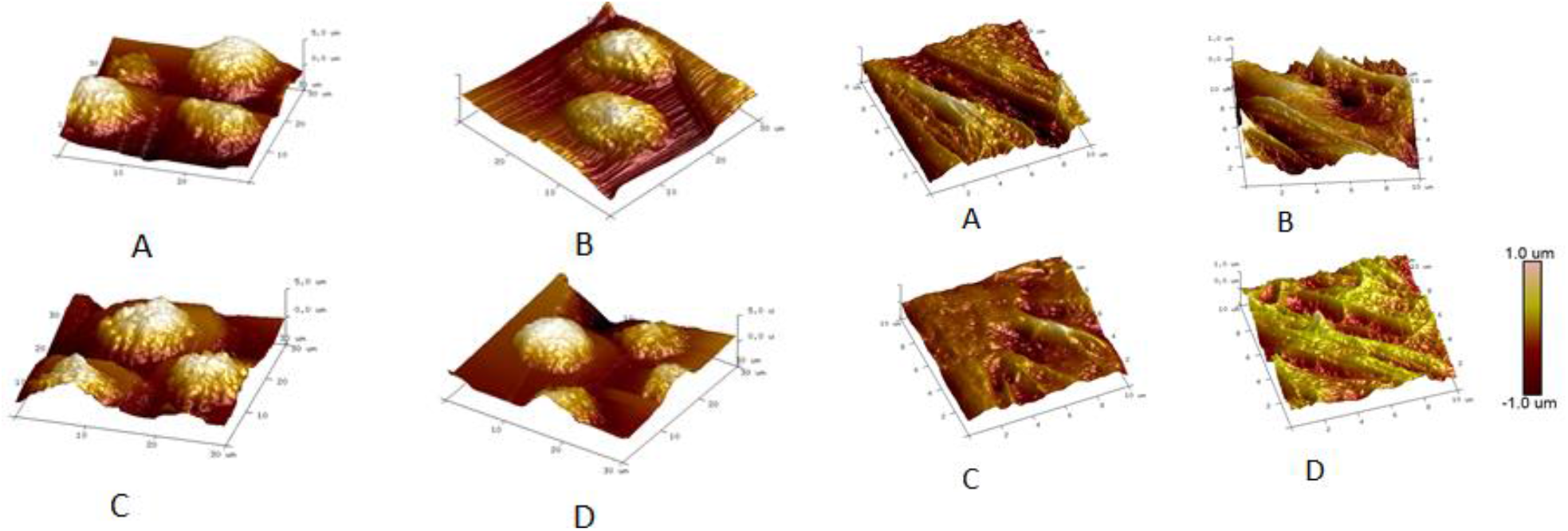
M1: Topography images of the articular chondrocytes measured by AFM (A: control, B: nicotine treatment, C: B treated by estrogen, D: estrogen treatment). Scan size: 30μm*30μm M2: Topography images of the articular chondrocytes adding estrogen receptor antagonist measured by AFM (A: control, B: nicotine treatment, C: B treated by estrogen, D: estrogen treatment). Scan size: 10μm*10μm in vitro

## 4. Discussion

### 4.1 Effect of nicotine and estrogen on the lubrication of articular cartilage

This study sought to elucidate the effect of nicotine and estrogen on the articular cartilage through difference of lubrication in vitro. Results showed that (1) nicotine significantly increased friction coefficient μ of cartilage for both blank (M1) and estrogen <u>receptor antagonist</u> (M2) articular cartilage in vitro. (2) After nicotine treatment, estrogen reduced μ significantly again, though the μ did not return that of control. (3) Adhesion force of M1 and M2 samples increased significantly after nicotine treatment and then reduced by estrogen treatment. The variation trend is similar to (1).

Decreased boundary lubrication performance is related to degenerative joint diseases, such as osteoarthritis (OA)(Desrochers et al., 2013). Experiments have found that nicotine stimulation leads to a decrease in the friction coefficient of articular cartilage, indicating that nicotine stimulation leads to a decrease in the lubrication efficiency of the articular cartilage surface, and the possibility of its wear is higher. It shows that for arthritic cartilage, the stimulation of nicotine will aggravate the abrasion of the cartilage surface. At the same time, it was found that the lack of estrogen also led to an increase in the friction coefficient of articular cartilage and cell surface. Nicotine stimulation reduces the friction properties of cartilage and its cell surface, but estrogen reverses this effect, which is also consistent with reports in the literature (Son and Chun, 2018; Thornton et al., 2015).

### 4.2 Changes in secretion of PRG4 for articular cartilage

These findings suggest that the proteinaceous component of the articular cartilage surface is changed due to nicotine and estrogen treatment. Reports showed that PRG4 is proposed as the key boundary lubricant of articular cartilage and the lack of PRG4 is related to the early onset of OA. In vitro cell confocal experiments can show that for M1 and M4, the content of PRG4 is significantly reduced after nicotine treatment, and its content is increased under estrogen treatment. This trend is consistent with the changing trend of friction capacity is just the opposite. It can be seen that nicotine and estrogen will affect the secretion of PRG4 by articular cartilage cells, and then affect the friction properties of the surface.

These results may be attributed to the expression levels of lubricants on the surface of the articular cartilage as evidenced by the immunohistochemistry results. After nicotine treatment, The level of inflammatory factors(TNF-α, IL-1β) was detected in our previous work(Peng et al., 2019). The increase in μ after nicotine treatment indicates a loss of the boundary lubricant, with PGR4 as the strongest candidate. The SZ accounts for the expression of the zone-specific secreted glycoprotein proteoglycan4(Grogan et al., 2013; Sakata et al., 2015). In recent years, research has found that PGR4 may slow or treat the progression of osteoarthritis(Ruan et al., 2013) and (Prg4)-null exhibit irreversible osteoarthritis-like changes. PRG4 is important for joint lubrication in reducing friction and wear(Chan et al., 2010; Coles et al., 2010a). Microscale surface friction of articular cartilage in early osteoarthritis is increased and proteoglycan4 expression protects against the development of osteoarthritis(Ruan et al., 2013). And increased friction is associated with loss of superficial zone chondrocytes that secreted lubricin(Coles et al., 2010b). The results of the present investigation, the decrease of lubrication describes the evidence that nicotine treatment destroys the secretion of PGR4 in the articular cartilage. Furthermore, confocal results show that the content of PRG4 decreased significantly(Fig.3.1) for both samples of blank and adding estrogen receptor antagonist. These demonstrate that high concentration of nicotine treatment would reduce the secretion of PGR4 from the articular cartilage.

**Fig.3.**
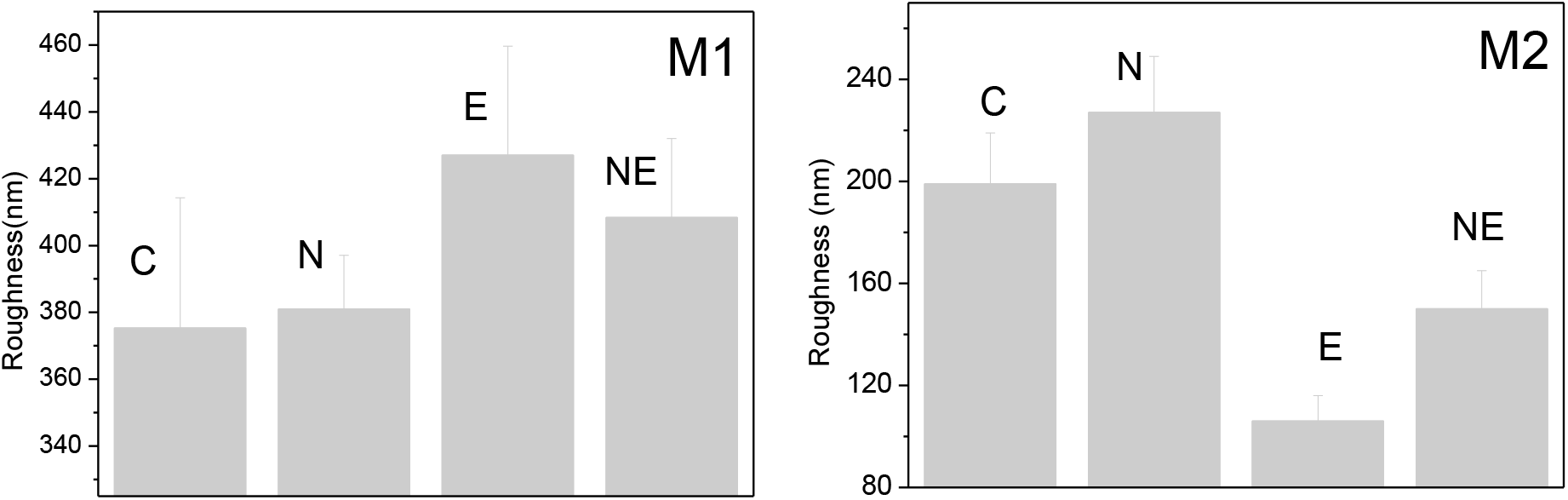
The RMS roughness of articular chondrocytes surfaces treatment with ((M1 for normal chondrocytes, M2 for chondrocytes of adding estrogen receptor antagonist, C: control; N: nicotine treatment; E: only estrogen treatment; NE: estrogen treating after nicotine treatment) in vitro

**Fig.4.**
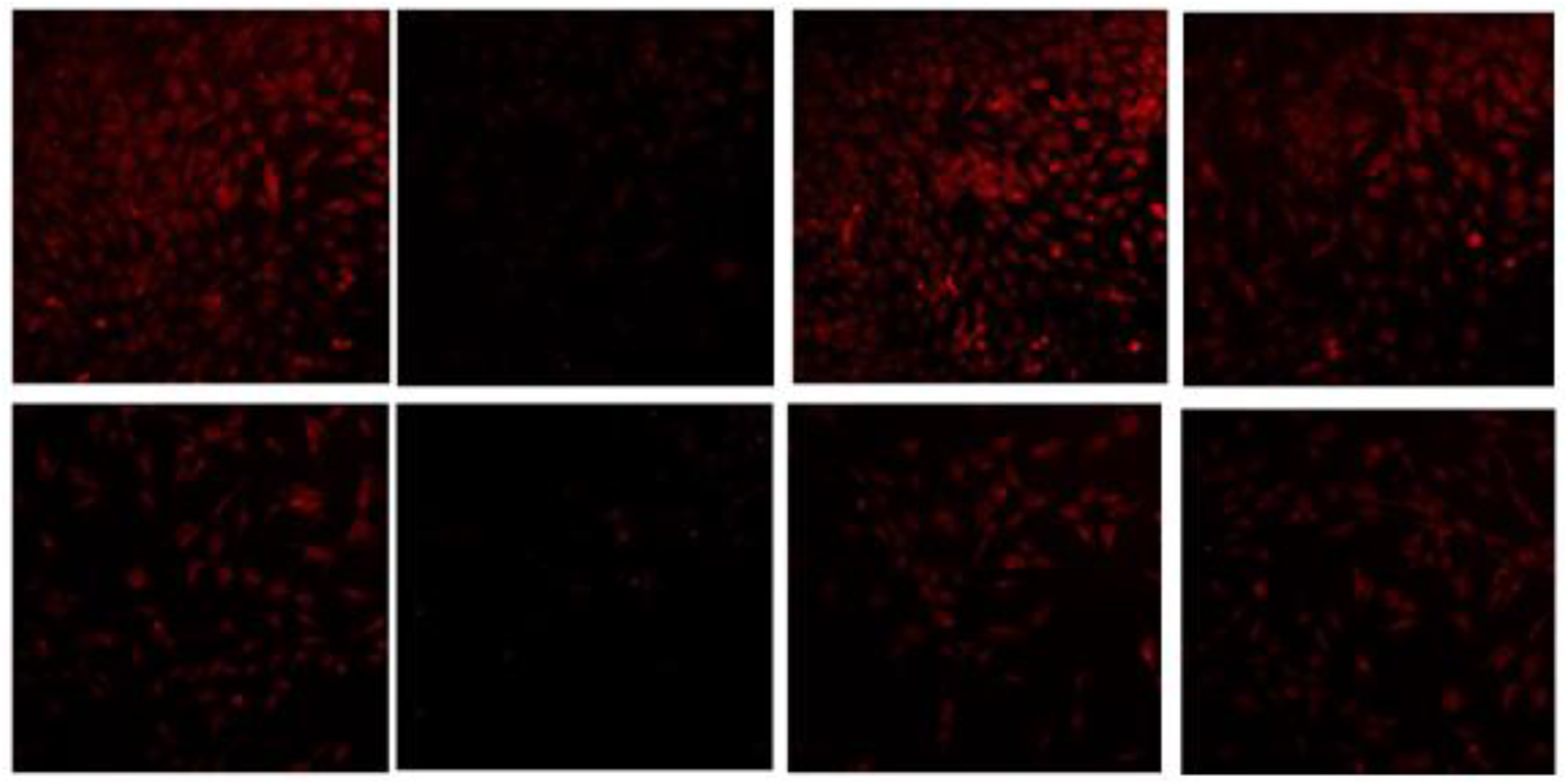
Adhesion force measured by AFM increased significantly after nicotine treatment and recovered a lot after estrogen treatment both for M1 and M2 in vitro ((M1 for normal chondrocytes, M2 for chondrocytes of adding estrogen receptor antagonist, C: control; N: nicotine treatment; E:only estrogen treatment; NE: estrogen treating after nicotine treatment)

The estrogen was used to and the friction on the cell was measured. The μ and PRG4 content were both recovered a lot which indicates the protective effect of the estrogen on the lubrication as verified in this study and previously(Peng et al., 2019). There are seldom reports to state relation between estrogen and friction of the articular cartilage, though, the positive effect of the estrogen were studied through different pathway. Physiological concentrations of estrogen prevented the injury-related cell death and reduced the GAG release significantly in a receptor-mediated manner(Imgenberg et al., 2013). Users of oestrogen replacement therapy had more knee cartilage than non-users indicated that ERT prevents loss of knee articular(Wluka et al., 2001). This study confirms the therapeutic effect of estrogen on PRG4 deficiency at the molecular level, however the physiological concentrations of estrogen did not have a significant effect on the friction and viscosity of the control group.

The lubrication mechanism of the articular cartilage was studied by AFM(Raj et al., 2017; Seror et al., 2011; Wang et al., 2013). And the research showed that in different stage OA, the cartilage superficial layer showed a higher μ the normal cartilage surface in PBS(Desrochers et al., 2013; Park et al., 2014). Friction coefficient changes did not correspond to surface roughness changes, consistent with previous report(Chan et al., 2010; Coles et al., 2010a); However adhesion increased significantly after nicotine treatment.

### 4.3 Adhesion between the AFM tip and the articular cartilage

Adhesion between the AFM tip and the cartilage surface was significant and directly related to the friction response. The higher friction and adhesion of nicotine treatment cartilage than control also correlated with the lower expression levels of PRG4. The higher adhesion of nicotine treatment cartilage may also be attributed to the decreasing of anti-adhesive and chondroprotective properties of SZP, which prevent synoviocyte overgrowth and cartilage-cartilage adhesion(Chan et al., 2011; Englert et al., 2005; Greene et al., 2015; Rhee et al., 2005). Boundary lubricant, (also known as lubricin and PRG4), was prevented to secrete due to nicotine treatment. And then estrogen treatment may stimulate the secretion of the PRG4 back again and lower adhesion at the surface of the cartilage, reduce articular cartilage friction.

The results of the present study demonstrate a persistent dependence of the friction coefficient of articular cartilage on nicotine and estrogen treatment by effect secretion of the PRG4.

Nicotine treated M1 and M2 demonstrated significantly higher adhesion forces than controls (f_C_=0.08nN vs f_N_=0.22nN for M1 and f_C_=0.18nN vs f_N_=1.22nN for M2 P< 0.0001), and after that estrogen treatment resulted in significantly lower adhesion forces for both M1 (P < 0.0001) and M2 (P < 0.0005)(Fig.5) But differences in adhesion between control and estrogen treatment samples were insignificant for both M1 and M2. The variation trend of adhesion force, demonstrating a pattern similar to that of the friction coefficient was also observed.

**Fig.5.**
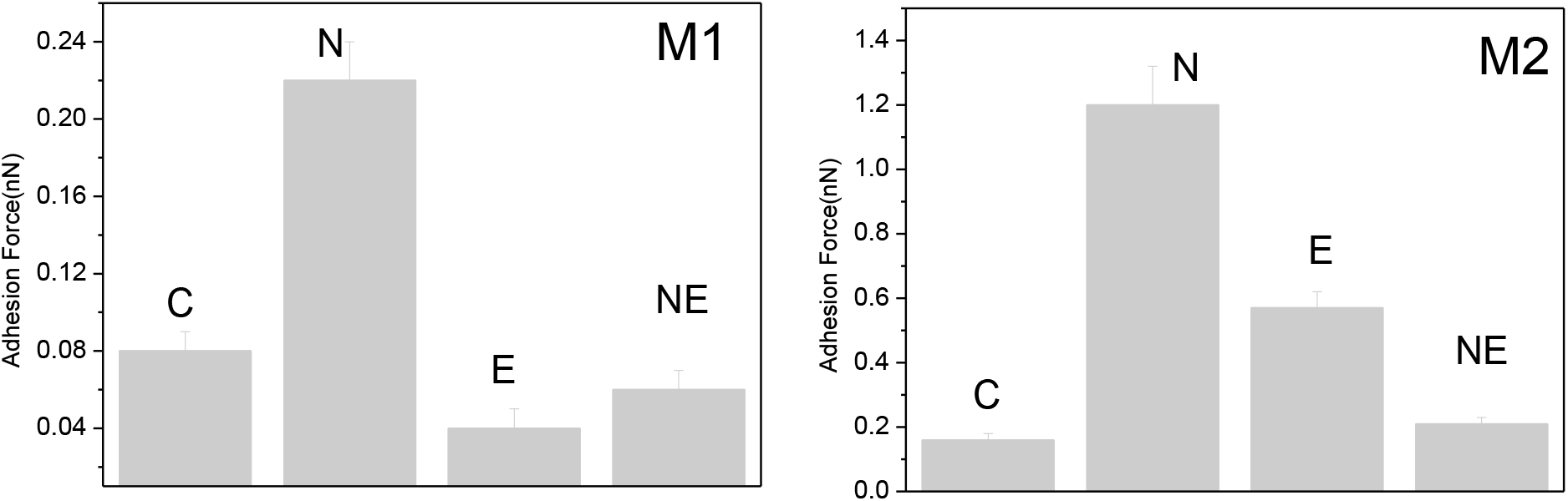
The proteoglycan content stained with safranin O (red) decreased after trypsin treatment, particularly in the uppermost layer of the superficial zone. The collagen content stained with light green.

## 5. Conclusion

The friction performance and adhesion force on the surface of articular chondrocytes were investigated using atomic force microscope. The articular chondrocytes and cartilage were stimulated by nicotine and then treated by estrogen, the results showed that nicotine down-regulates the friction coefficient and secretion of PRG4 significantly and then the estrogen increase them again. The adhesion forces also showed the same trend due to the content of anti-adhesive PRG4. This study demonstrated that the present concentration nicotine has a negative effect on the articular cartilage and the estrogen has a better protecting effect. This may provide a potential guide for OA prevention and treatment.

